# Invertebrate DNA methylation and gene regulation

**DOI:** 10.1101/2021.11.19.469267

**Authors:** Groves Dixon, Mikhail Matz

**Affiliations:** Department of Integrative Biology, University of Texas at Austin

## Abstract

As human activity alters the planet, there is a pressing need to understand how organisms adapt to environmental change. Of growing interest in this area is the role of epigenetic modifications, such as DNA methylation, in tailoring gene expression to fit novel conditions. Here, we reanalyzed nine invertebrate (Anthozoa and Hexapoda) datasets to validate a key prediction of this hypothesis: changes in DNA methylation in response to some condition correlate with changes in gene expression. While we detected both differential methylation and differential expression, there was no simple relationship between these differences. Hence, if changes in DNA methylation regulate invertebrate transcription, the mechanism does not follow a simple linear relationship.

## Introduction

The regulatory function of DNA methylation in invertebrates, if any, remains unclear. Methylation of promoters, which is important for mitotically heritable gene silencing in vertebrates, has not been observed in invertebrates (Zemach et al. 2010; de Mendoza et al. 2019; Xu et al. 2019). In contrast to promoter methylation, gene body methylation (gbM) is positively correlated with gene expression in both invertebrates and plants. There is little evidence however, that it directly regulates transcription. For instance, several studies failed to detect substantial differences in gbM across invertebrate cell types or developmental stages (Gatzmann et al. 2018; Harris et al. 2019; de Mendoza et al. 2019) despite profound transcriptome differences (although see Liew et al. (2018). Conversely, removal of gbM by knockdown of DNMT1 enzyme did not significantly alter gene expression in a milkweed bug (Adam J. Bewick et al. 2019). Similar results have been observed in plants (Bewick et al. 2016; Adam J Bewick et al. 2019; Choi et al. 2020). Together, these studies indicate that changes in gbM are neither necessary nor sufficient to induce changes in transcription.

Here, we re-analyzed publicly available methylomic and RNA-seq data from three Anthozoa and six Hexapoda studies to evaluate relationships between invertebrate DNA methylation and transcription. For each study, we contrast methylation- and transcriptional differences between two experimental conditions. The Anthozoan studies contrasted polyp types in the coral *Acropora millepora* (Dixon and Matz 2020), pH treatments in the coral *Stylophora pistillata* (Liew et al. 2018), and symbiotic state in the sea anemone *Exaiptasia pallida* (Li et al. 2018). Hexapoda studies included different reproductive states in ants (*Ooceraea biroi*)(Libbrecht et al. 2016), bumblebees (*Bombus terrestris*) (Marshall et al. 2019), and termites (*Zootermopsis nevadensis*) (Glastad et al. 2016), different subcastes in honeybee (*Apis mellifera*)(Herb et al. 2012), differences in maternal care in carpenter bee (Arsenault et al. 2018), and different diapause states in silkworm (*Bombyx mori*)(Li et al. 2020). Using these diverse datasets, we first confirm previous findings that baseline gbM levels are bimodally distributed across coding genes, are positively associated with baseline transcription level, and are negatively associated with transcriptional differences between conditions. We next examine the hypothesis that changes in gbM and promoter methylation between conditions correlate with changes in transcription, but find no support for this prediction.

## Methods

### Previously published datasets

Previously published WGBS and RNA-seq datasets from invertebrate species are shown in Figure 1. The criteria for selecting these projects were: 1) the project focused on an invertebrate species 2) the project included at least two conditions, such as environmental exposure, or caste. 3) the project characterized DNA methylation using Whole Genome Bisulfite Sequencing (WGBS) 4) the project characterized transcription using RNA-seq 5) reads were available on the NCBI SRA database. Experimental methods from some projects allowed for multiple comparisons, however for simplicity, we focused on contrasts that seemed likely to induce the greatest epigenetic change. The comparisons we made are as follows. For the anemone *Exaiptasia pallida* (Li et al. 2018), we compared aposymbiotic (N=6) to symbiotic (N=6) individuals. For the smooth cauliflower coral *Stylophora pistillata* (Liew et al. 2018), we compared only the most extreme pH treatment (pH 7.2; N=3) to controls (pH 8.0; N=3). For silkworm *Bombyx mori* (Li et al. 2020), we compared diapause terminated (N=3) to diapause destined (N=3) eggs. For the termite *Zootermopsis nevadensis nuttingi* (Glastad et al. 2016), we compared winged reproductive alates of both sexes (N=4) to larval instars (workers) of both sexes (N=4). For the small carpenter bee *Ceratina calcarata* (Arsenault et al. 2018) we compared newly eclosed adults that developed without maternal care (N=3) to those that received maternal care (N=3). For bumblebee *Bombus terrestris* (Marshall et al. 2019), we compared reproductive (N=3) to sterile castes (N=3). For honeybee *Apis mellifera* (Herb et al. 2012), we compared nurse subcastes (N=6) to worker subcastes (N=6). For the clonal raider ant *Ooceraea biroi* (Libbrecht et al. 2016), we compared individuals in the reproductive phase (N=4) to those in brood care phase (N=4).

**Figure 1:**
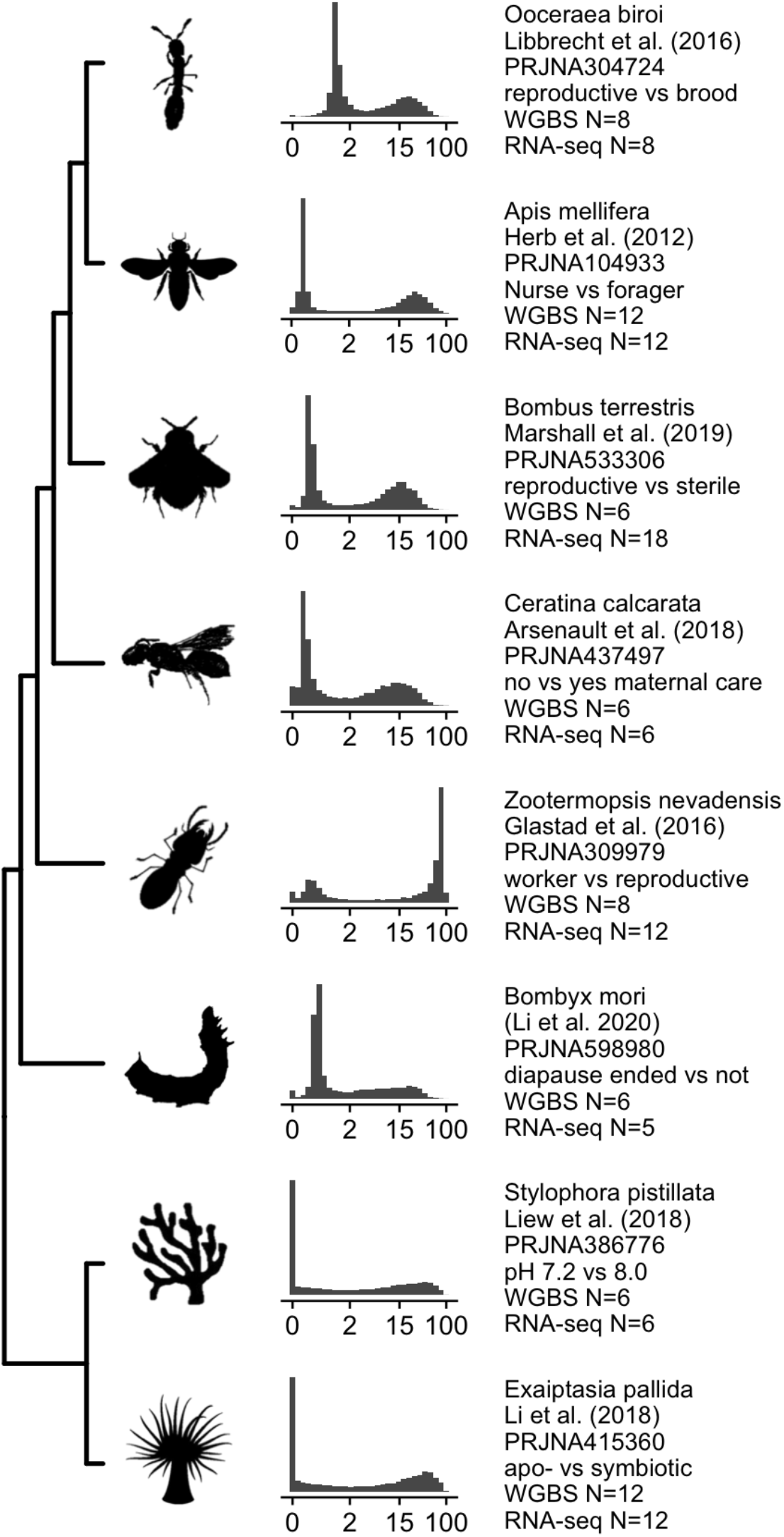
Overview of species covered by previously published datasets and the distribution of gbM levels in each. X-axes for histograms show percent methylation (summed across all CpG sites within the gene) on the log_2_ scale. Text reports the species, reference, NCBI Bioproject accession, the treatment groups compared here, the Whole Genome Bisulfite sample size, and the RNA-seq sample size for each study.

### WGBS data processing

Raw reads were trimmed and quality filtered using cutadapt, simultaneously trimming low-quality bases from the 3’ end (-q 30) and removing reads below 50 bp in length (-m 50) (Martin 2011). Trimmed reads for each dataset were mapped to the appropriate reference genome (Table S1; Xia et al. 2008; Baumgarten et al. 2015; Sadd et al. 2015; Rehan et al. 2016; Voolstra et al. 2017; McKenzie and Kronauer 2018; Fuller et al. 2020) using Bismark v0.17.0 (Krueger and Andrews 2011) with adjusted mapping parameters (--score_min L,0,-0.6). Reads from Dixon and Matz (2020) were mapped using -- non_directional mode as recommended by the Pico Methyl-Seq Library Prep Kit manual. PCR duplicates were removed from the Bismark alignment files using the deduplicate_bismark command. Methylation levels were extracted from the alignments using bismark_methylation_extractor with the -- merge_non_CpG, --comprehensive, and --cytosine_report arguments. Detailed steps used to process the WGBS reads are available on the git repository (Dixon 2020).

### RNA-seq data processing

Raw reads were trimmed and quality filtered using cutadapt, simultaneously trimming low-quality bases from the 3’ end (-q 30) and removing reads below 50 bp in length (-m 50)(Martin 2011). Trimmed reads for each dataset were mapped to the appropriate reference genome (Table S1) using Bowtie2 using the --local argument (Langmead and Salzberg 2012). PCR duplicates were removed using MarkDuplicates from Picard Toolkit (Broad Institute 2019). Sorting and conversion from sam files was performed using Samtools (Li et al. 2009). The reads mapping to annotated gene boundaries were counted using FeatureCounts (Liao et al. 2014). Detailed steps used to process the RNA reads are available in the git repository (Dixon 2020).

### Measuring gbM level

Based on previous findings that different measures of gbM were highly similar (Dixon and Matz 2020), we reported gbM level as the percent methylation rate on the log_2_ scale. Here the percent methylation rate for a gene is the ratio of the total number of methylated read counts to read all counts summed across all CpG sites within the bounds of the gene. To allow plotting on the log scale, zero values were assigned to the lowest non-zero value for each project. As methylation sometimes occurs preferentially on invertebrate exons, we also calculated gbM based only on exons. Focusing the analysis on only exons resulted in clearer gbM class separation and stronger associations with gene expression for the *Acropora millepora* dataset, but not for the other datasets. For this reason, we separately reported gbM based on exons for *A. millepora*, but entire genes for the rest of the studies. Following previous studies (Dixon et al. 2016; Dixon et al. 2018; Dixon and Matz 2020), in the case of MBD-seq we report gbM as the log_2_ fold difference between the captured and unbound fractions generated during library preparation. gbM level based on mdRAD was computed as Fragments per Kilobase of gene length per Million reads (FPKM) on the log_2_ scale. Analyses of differential methylation based on bisulfite sequencing data were done using MethylKit package (Akalin et al. 2012).

### Relationships between gbM and mRNA level

For each dataset, we tested for expected relationships between gbM and mRNA expression patterns. For our dataset, generated using Tag-seq (Meyer et al. 2011), we calculated mean mRNA level by averaging the regularized counts generated using the rlog function in DESeq2 across all samples (Love et al. 2014). For the other datasets, which used standard RNA-seq, we calculated mean mRNA level as FPKM averaged across all samples. Differences in mRNA abundance between groups were calculated using DESeq2 (Love et al. 2014). For our dataset, this analysis was performed including colony identity (genotype) as a factor to control for genetic effects. For simplicity, models for differential expression for the published datasets included only the treatment groups indicated in Figure 1 (we did not include additional factors, for instance, sex or colony identity). Differences between groups are reported as log_2_ fold differences.

## Results and Discussion

### Confirming previous relationships between gbM and transcription

We first sought to corroborate previous findings on the distribution of gbM, and its relationship to gene expression patterns. Using three different methylation assays, we confirmed that gbM in *A. millepora* shows a characteristic bimodal distribution, separating genes into methylated and unmethylated classes (Figure 2 A-C). We then confirmed that gbM level is associated with average mRNA abundance (Figure 2 D-F), and negatively associated with differential expression between polyp types (Figure 2 G-I). Hence, regardless of the method used to measure methylation, gbM shows the expected distribution and associations with gene expression in *A. millepora*.

**Figure 2:**
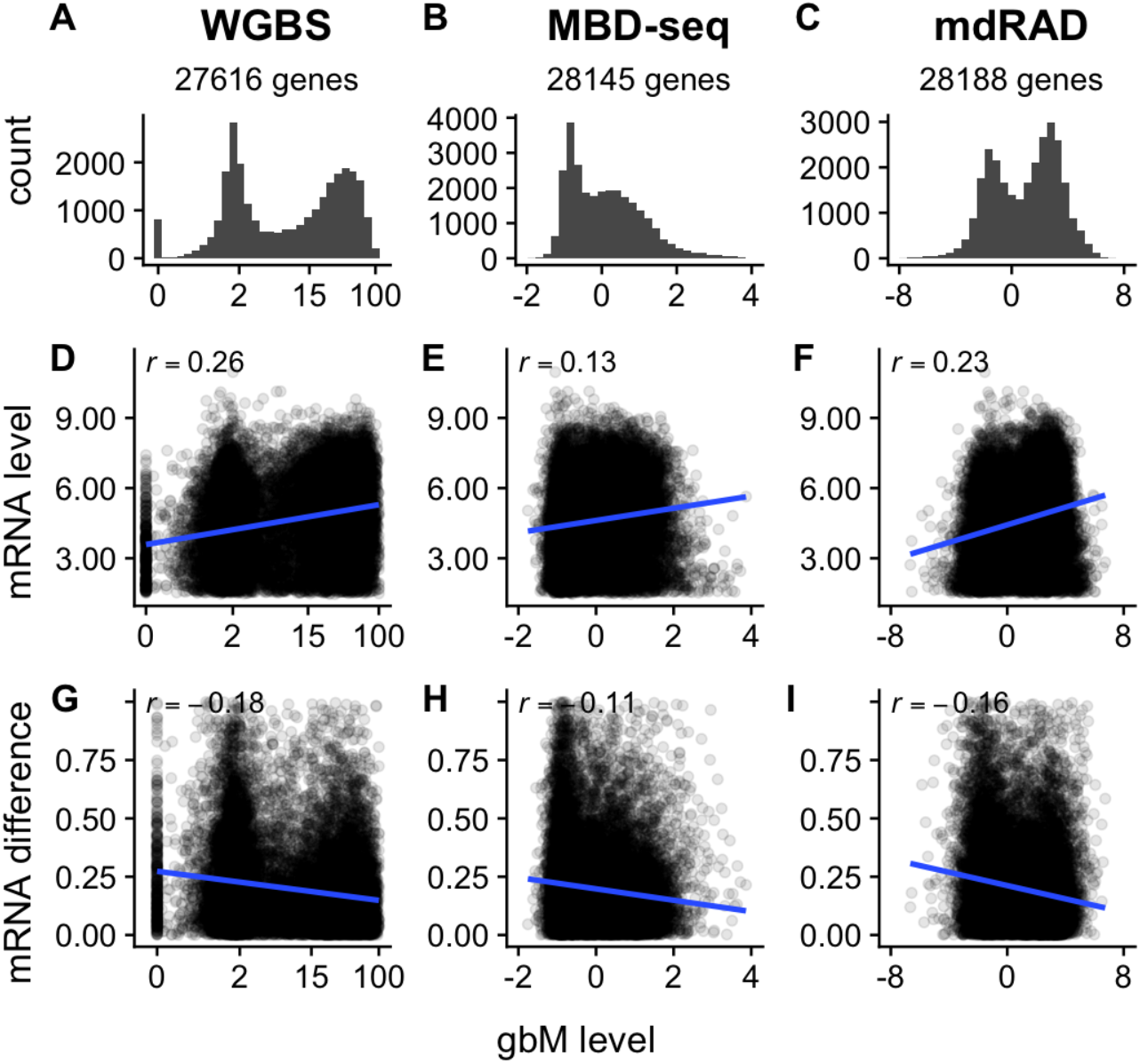
Associations between gbM level and gene expression patterns in *A. millepora* based on three different methylation assays. (A-C) Distribution of gbM levels (D-F) Relationship between gbM level and mRNA abundance (G-I) Relationship between gbM level and differential mRNA expression between axial and radial polyps. X-axes for the assays are as follows. Whole Genome Bisulfite Sequencing (WGBS): percent methylation (summed across all CpG sites within exons of each gene) on the log_2_ scale; Methylation Binding Domain Sequencing (MBD-seq): the log_2_ fold difference in fold coverage between the captured and unbound fractions (see methods); methylation dependent RAD-seq (mdRAD): fragments per kilobase of gene sequence per million reads on the log_2_ scale (log_2_(FPKM)).

We found similar results in other studies. While the relative sizes and means of the peaks varied by dataset, these were also bimodally distributed (Figure 1), and similarly associated with mRNA expression patterns (Figure 3). Relationships between average gbM and baseline mRNA expression tended to be stronger among the insects than the cnidarians, both for mean mRNA abundance and differential mRNA expression between groups.

**Figure 3:**
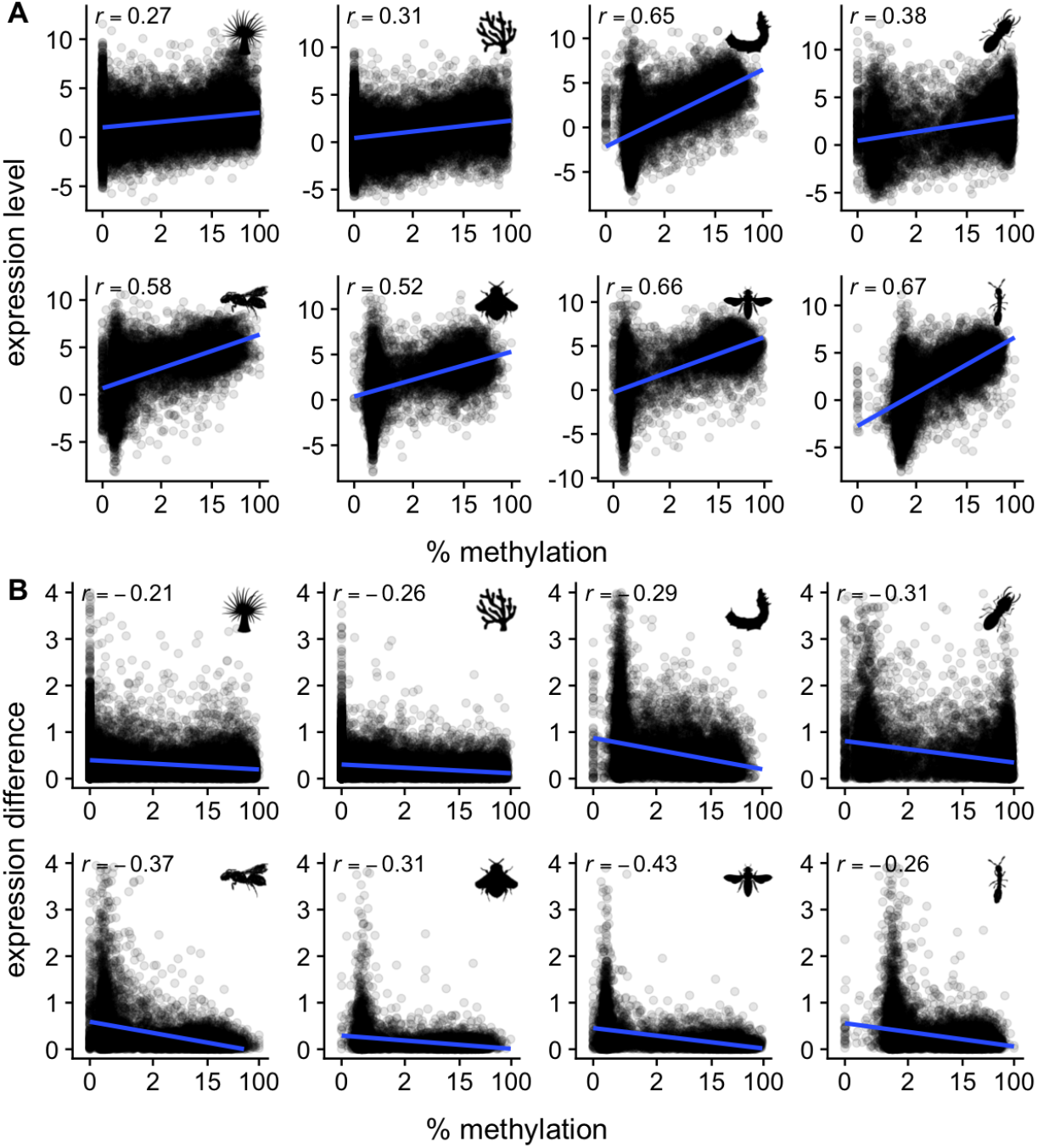
Associations between gbM level and gene expression patterns. (A) Relationship between gbM level and mRNA abundance (RNA-seq FPKM). (B) Relationship between gbM level and differential mRNA expression between treatment groups (Figure 1). X-axis shows percent methylation on the log_2_ scale. The correlation coefficient is given in the upper right hand of each plot.

### No correlation between changes in gbM and changes in transcription

As gbM is associated with elevated expression, a simple hypothesis is that increasing gbM increases transcription. Our re-analysis of gbM and mRNA differences between treatment groups shows that this is not the case. Using three different methylation datasets in *A. millepora*, we found that measurements of gbM differences between polyp types showed no consistent association with transcriptional differences (Figure S1). This was also the case for each of the other datasets (Figure S2). Hence, in Anthozoa and Hexapoda, gbM and transcription show no simple covariation between study conditions.

### No correlation between changes in promoter methylation and changes in transcription

As promoter methylation is associated with gene silencing in vertebrates, we tested whether changes in promoter methylation correlate with gene expression changes in invertebrates. Specifically, we tested whether differences in methylation in windows 1Kb upstream of genes between experimental groups predicted differences in mRNA levels. As with gbM, we found no reproducible relationship between differential promoter methylation and differential expression for *A. millepora* (Figure S3), or any of the other studies (Figure S4).

### Evidence for the “seesaw” hypothesis

As invertebrate coding genes are separated into methylated and unmethylated classes, we hypothesized that these class designations serve as a gene-regulatory signal, and that the methylated and unmethylated classes of genes undergo group-level changes in expression. In a previous study of *A. millepora* we observed inverse changes in both gbM and transcription depending on methylation class (the “seesaw hypothesis”, Dixon et al. (2018)) and sought to replicate those observations here. We first examined the new dataset from *A. millepora*. Between polyp types, we detected weak patterns similar to those previously reported. Specifically, based on MBD-seq and mdRAD, in axial compared to radial polyps the high gbM class increased in methylation level while the low-gbM class decreased in methylation. However, this pattern was not apparent for the WGBS dataset (Figure S5). Based on all three methylation assays, transcription of the low gbM class was somewhat upregulated, and the high gbM class somewhat downregulated (Figure S5). While this was consistent with our previous observations (Dixon et al. 2018), the correlations were too weak (correlation coefficients from -0.06 to - 0.1) to claim confident support of the hypothesis.

Among the other studies, there were three cases (silkworm, termite, and carpenter bee) of change in methylation based on gbM class, but in contrast to Dixon et al. (2018), in all these cases only the high-gbM class was changing (Figure S6). Also in contrast to Dixon et al. (2018), these class-level changes in methylation were not associated with class-level changes in transcription (Figure S7). Conversely, two studies demonstrated class-level changes in transcription but no corresponding class-level changes in gbM. In these two cases (honeybee and bumblebee), the low-gbM class was downregulated on average, and high-gbM class upregulated on average (Figure 4; Figure S7). In summary, while aspects of the “seesaw” hypothesis proposed by Dixon et al. (2018) were detected in several cases, it was not fully supported by any of the studies included here.

**Figure 4:**
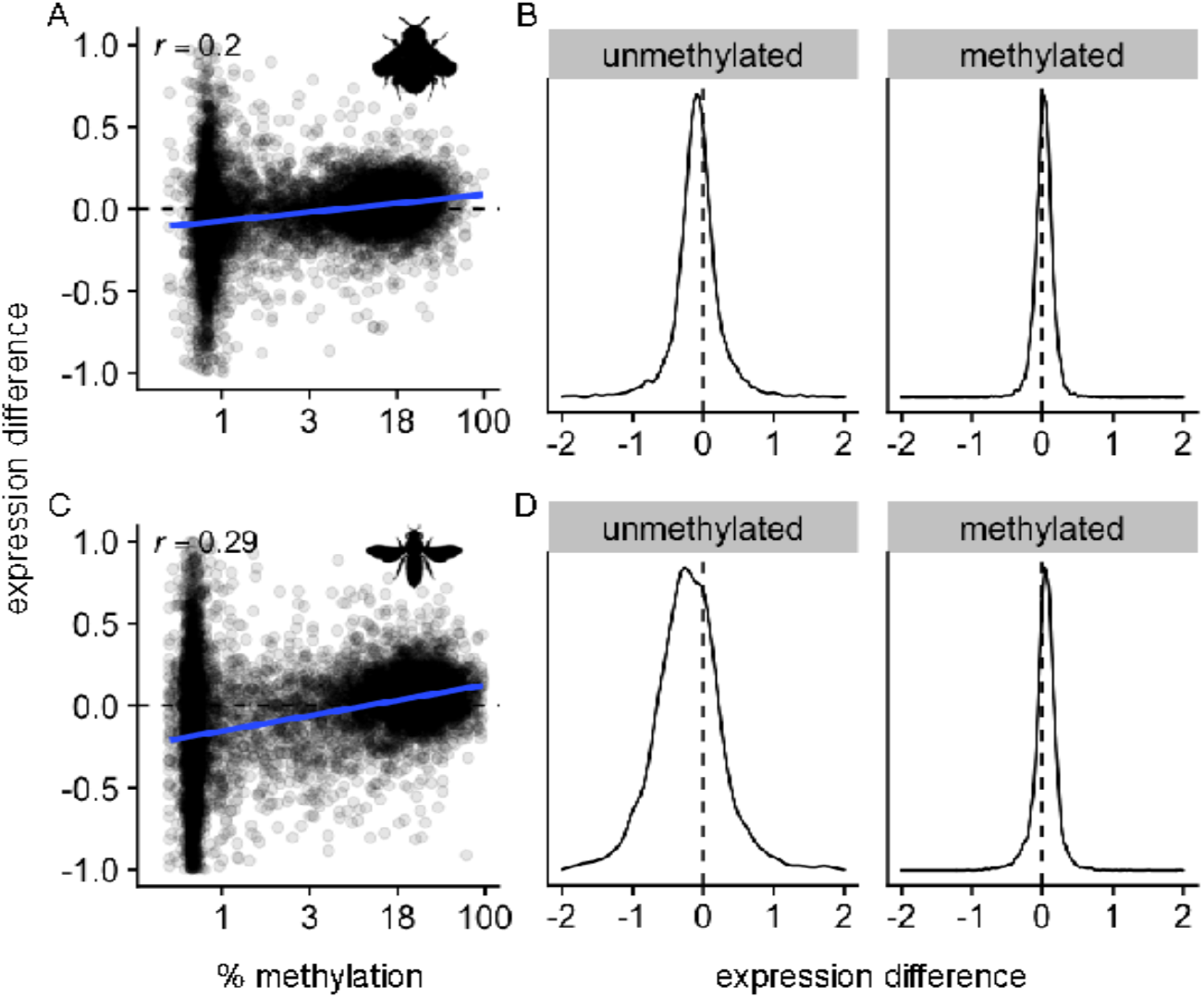
Two cases of inverse gbM class-level shifts in transcription. (A) Transcriptional differences plotted against gbM level for the bumblebee dataset (Marshal et al. 2019) contrasting reproductive vs sterile individuals. (B) Density plots of the log_2_ fold changes in transcription for the low-gbM (unmethylated) and high-gbM (methylated) classes. The unmethylated class tended to be downregulated and the methylated class tended to be upregulated. (C-D) Same plots contrasting nurse vs forager honeybee subcastes.

## Conclusions

Here we used published methylomic and transcriptomic data from Anthozoa and Hexapoda to examine how DNA methylation relates to transcriptional variation between different environmental and phenotypic conditions. We found that, as previously reported, gbM is bimodally distributed, and that higher gbM levels are associated with elevated transcription and less transcriptional variation between conditions. Differences in gbM between conditions showed no consistent linear association with differences in transcription. As there were often significant differences in both gbM and transcription (Figure S8), this indicates that changes in gbM alone are neither necessary nor sufficient to induce changes in transcription. Methylation differences between conditions 1 Kb upstream up the first exon also showed no association with differences in transcription.

In conclusion, if shifting methylation patterns regulate invertebrate transcription, the mechanism is more complex than can be captured by a simple linear relationship between these two variables. One possibility is that DNA methylation interacts with other epigenetic modifications (Bogan, Samuel et al. 2021), which must be included to accurately capture complex interrelationships relationship between methylation, other epigenetic modifications, and expression.

## Supplemental Material

**Table S1:** Reference genomes used in this study

| Species | Bioproject | Accession | Reference |
| --- | --- | --- | --- |
| *Exaiptasia pallida* | PRJNA261862 | GCA_001417965.1 | (Baumgarten et al. 2015) |
| *Acropora millepora* | PRJNA633778 | GCA_013753865.1 | (Fuller et al. 2020) |
| *Bombus terrestris* | PRJNA45869 | GCA_000214255.1 | (Sadd et al. 2015) |
| *Ceratina calcarata* | PRJNA299559 | GCA_001652005.1 | (Rehan et al. 2016) |
| *Ooceraea biroi* | PRJNA420369 | GCA_003672135.1 | (McKenzie and Kronauer 2018) |
| *Bombyx mori* | PRJDA20217 | GCF_000151625.1 | (Xia et al. 2008) |
| *Stylophora pistillata* | PRJDA20217 | GCA_002571385.1 | (Voolstra et al. 2017) |

**Figure S1:**
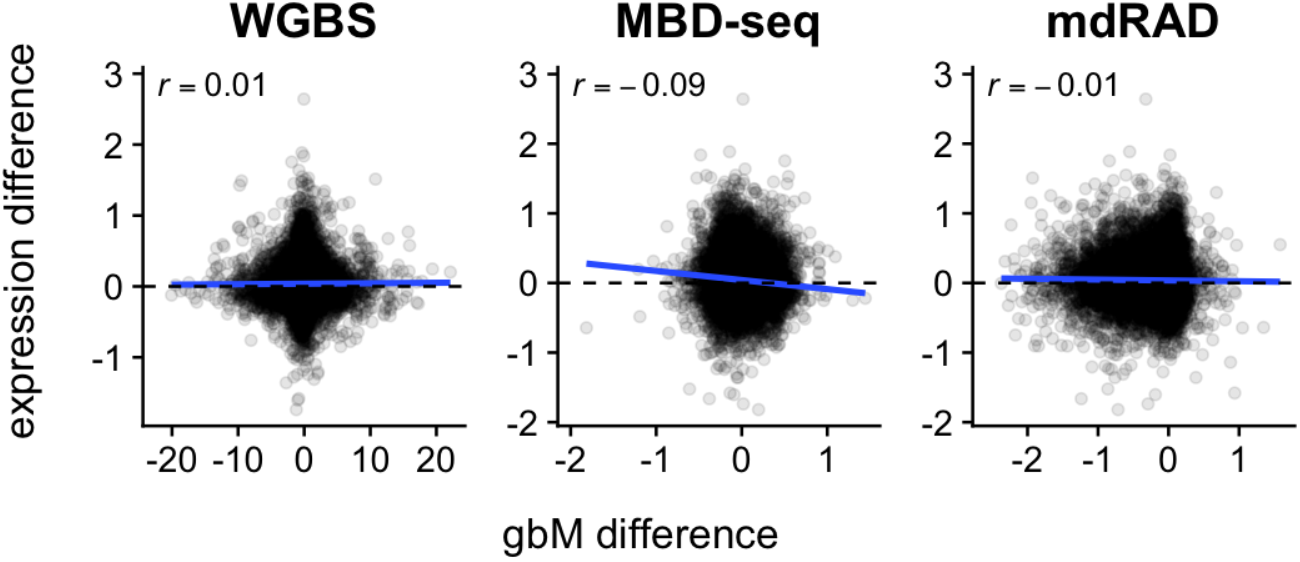
gbM and transcriptional differences between polyp types in *A. millepora* show no reproducible relationship. The title of each panel indicates the assay used to measure gbM differences. All axes are on the log_2_ scale.

**Figure S2:**
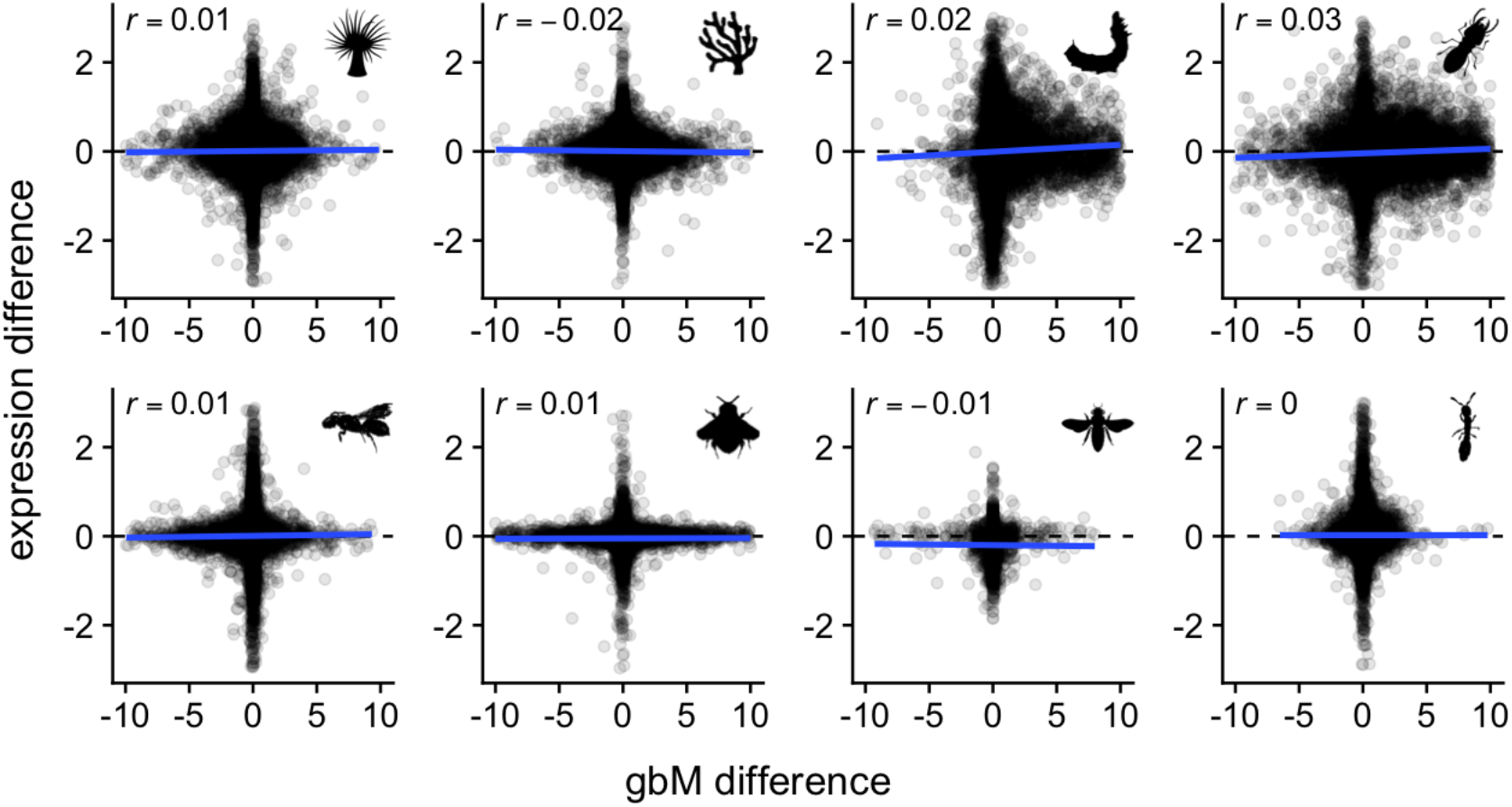
gbM and transcriptional differences between experimental conditions show little or no relationship. The contrasts for each study are given in Figure 1. All axes are on the log_2_ scale.

**Figure S3:**
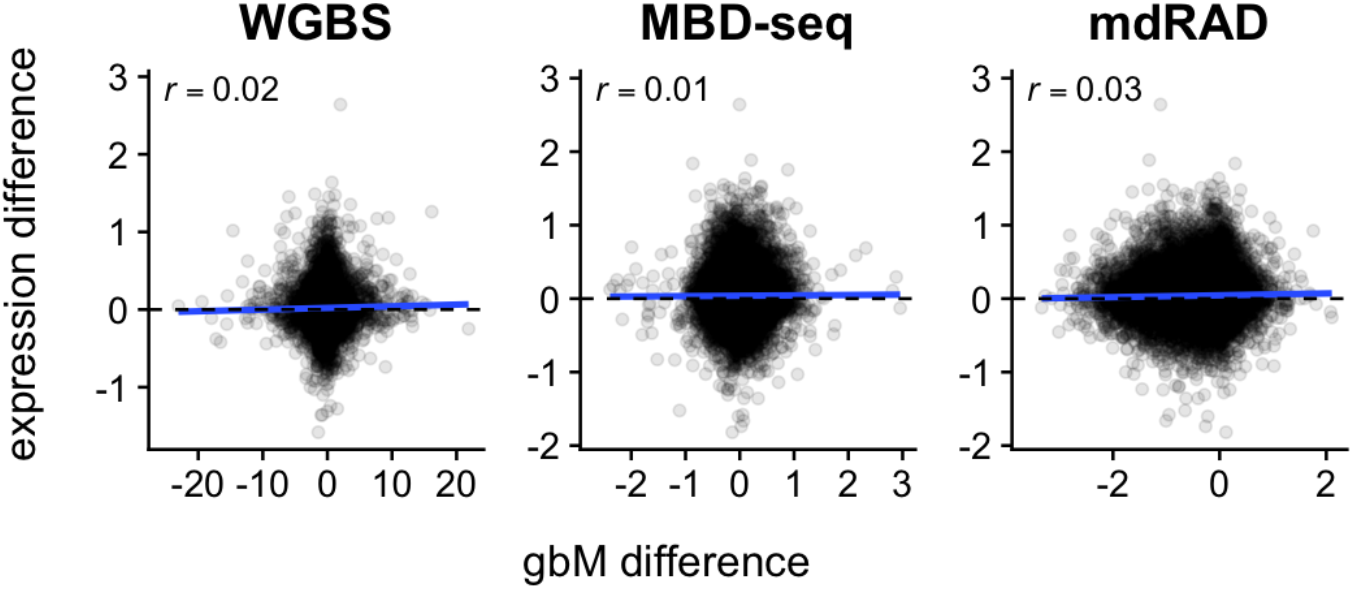
Promoter methylation and transcriptional differences between polyp types in *A. millepora* show no relationship.

**Figure S4:**
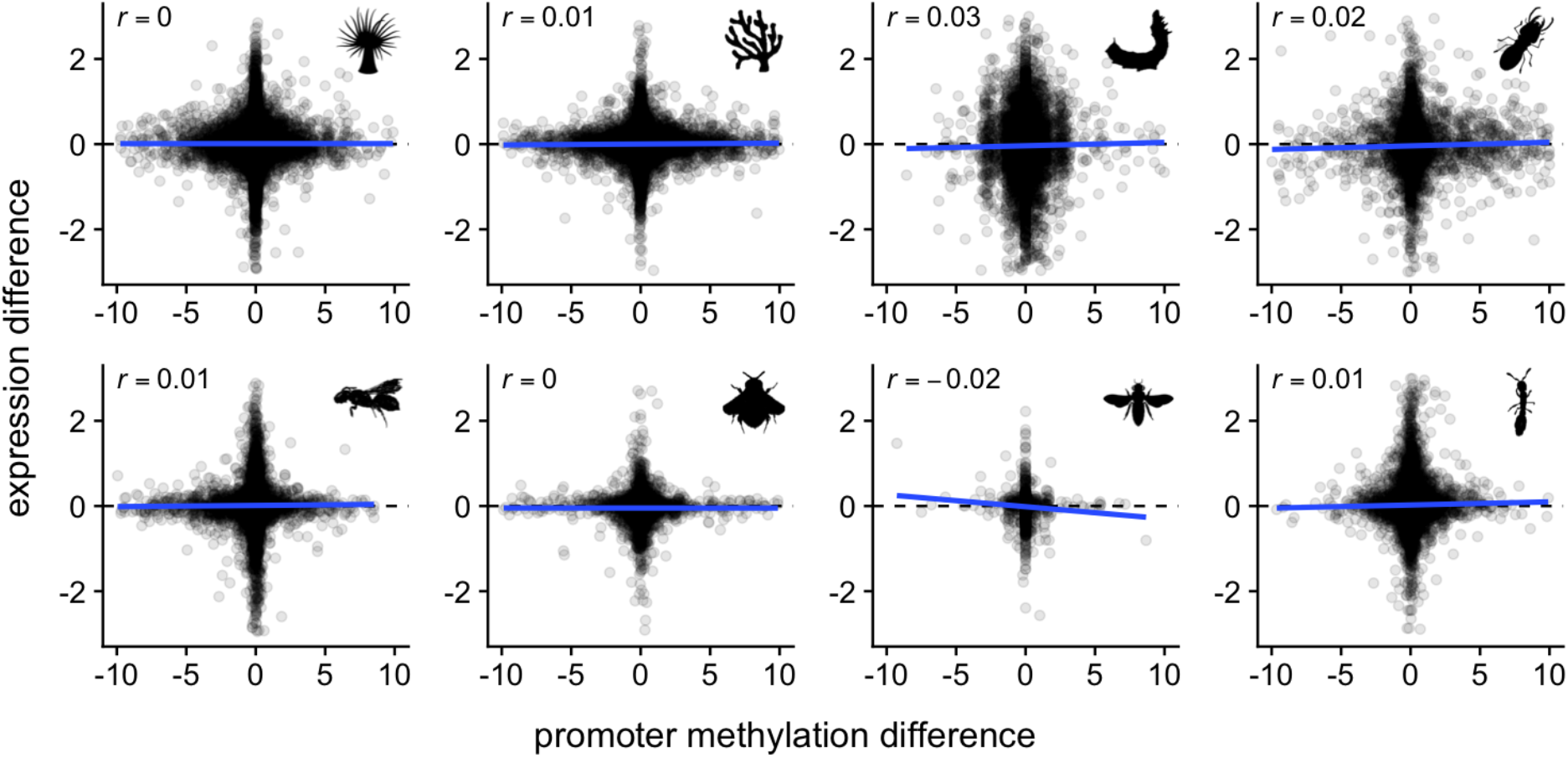
Promoter methylation and transcriptional differences between experimental conditions show no relationship.

**Figure S5:**
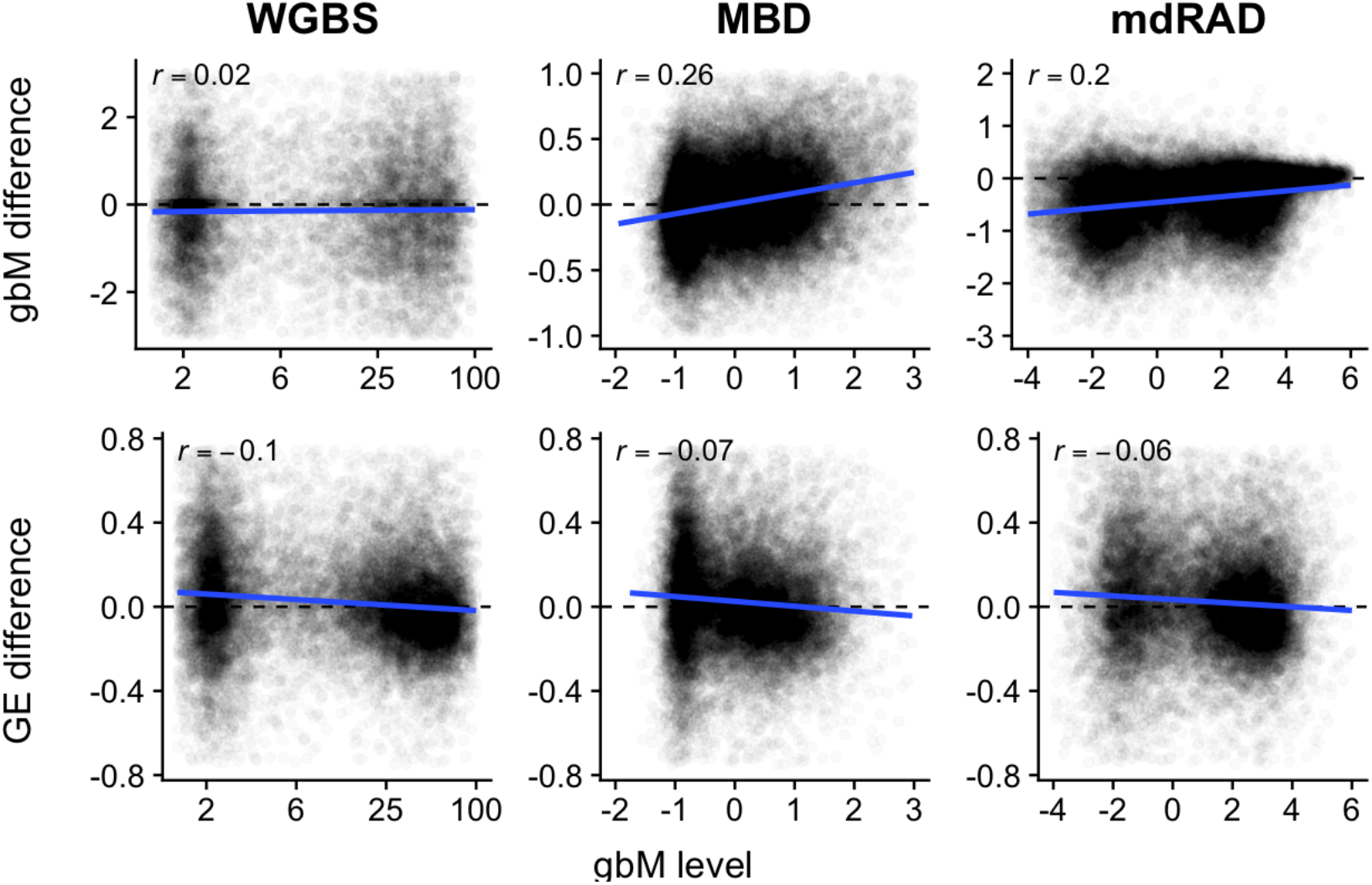
Relationships between differential gbM and gene expression between *A. millepora* polyp types and baseline gbM level. X axes show baseline gbM level. Y axes shows differential gbM between polyp types (top panels) and differential transcription between polyp types (bottom panels). Top panels: Differential gbM between polyp types was linked with baseline gbM level for the MBD-seq and mdRAD datasets, but not for WGBS. Bottom panels: For all three methylation datasets, the low gbM class of genes was somewhat upregulated in axial compared to radial polyps, and the high gbM class was somewhat downregulated.

**Figure S6:**
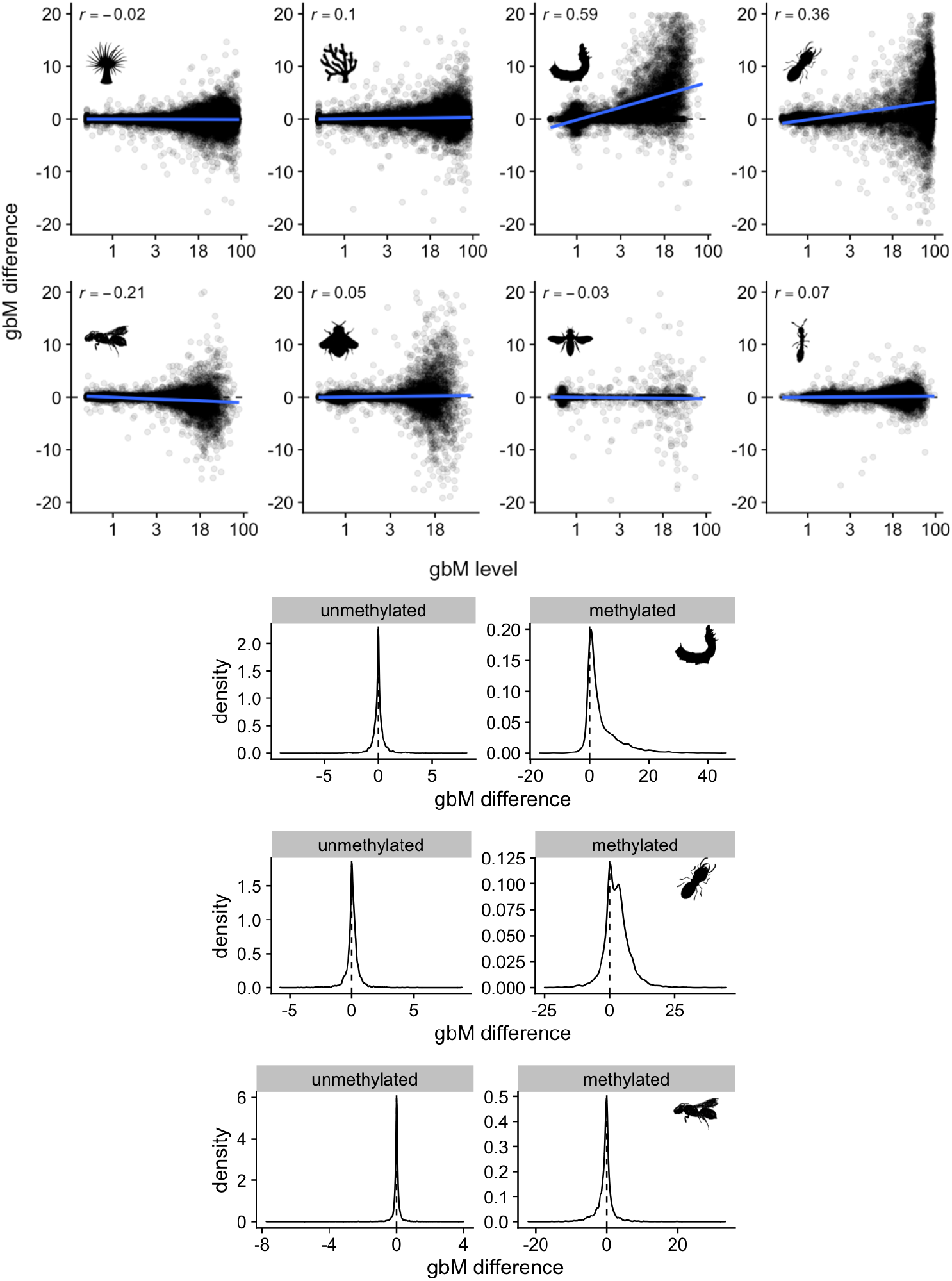
Scatterplots of gbM change against mean gbM level. Only three studies (Silkworm, Termite, and Carpenter bee) showed changes in methylation between conditions based on gbM class. Density plots of methylation change for the two classes show the change was specific to the high gbM class.

**Figure S7:**
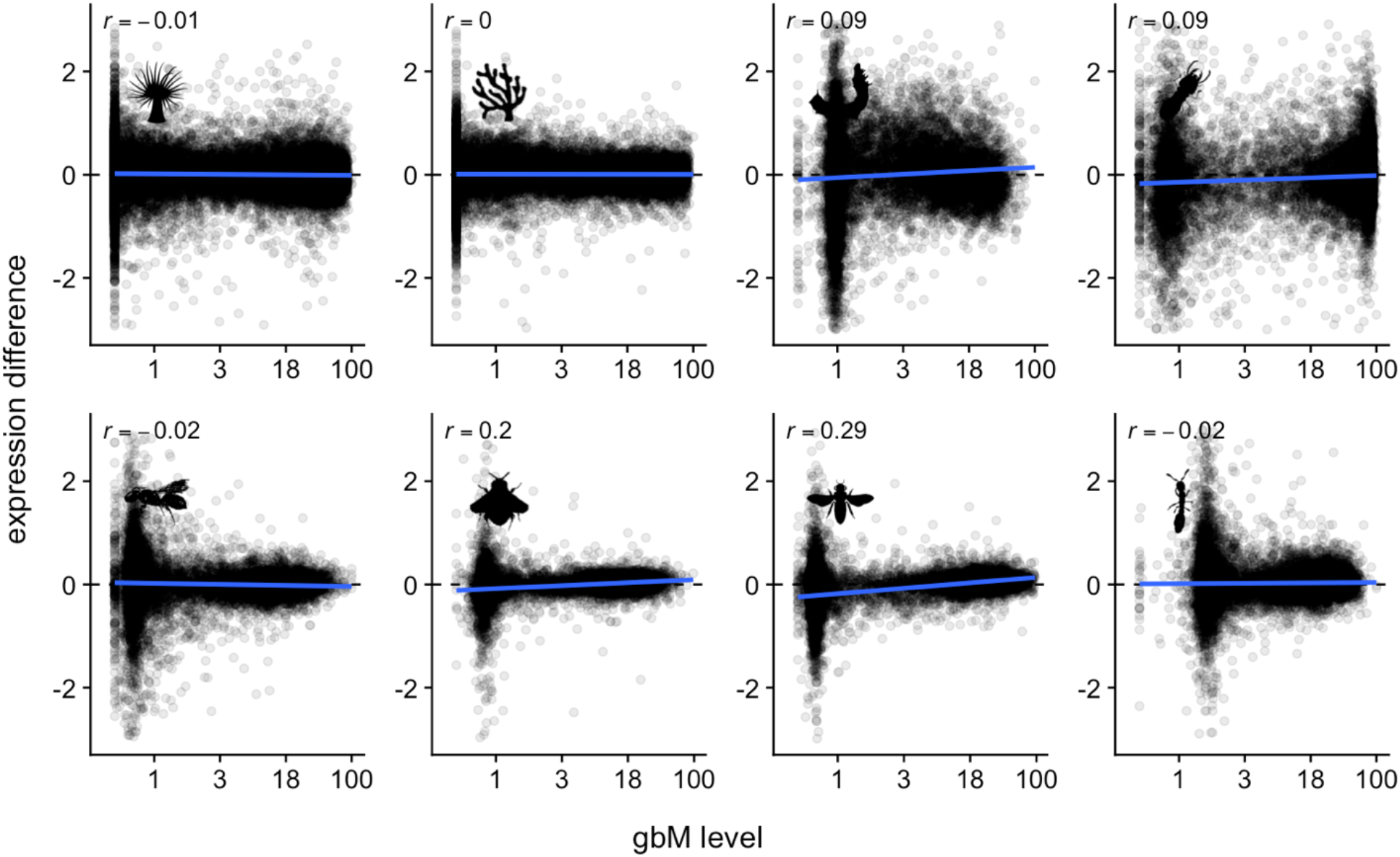
Scatterplots of transcriptional change against mean gbM level. Only two studies (bumblebee and honeybee) showed changes in transcription based on gbM class. These two are shown in greater detail in Figure 4. Note that these were distinct from the studies showing class-level changes in gbM (Figure S6).

**Figure S8:**
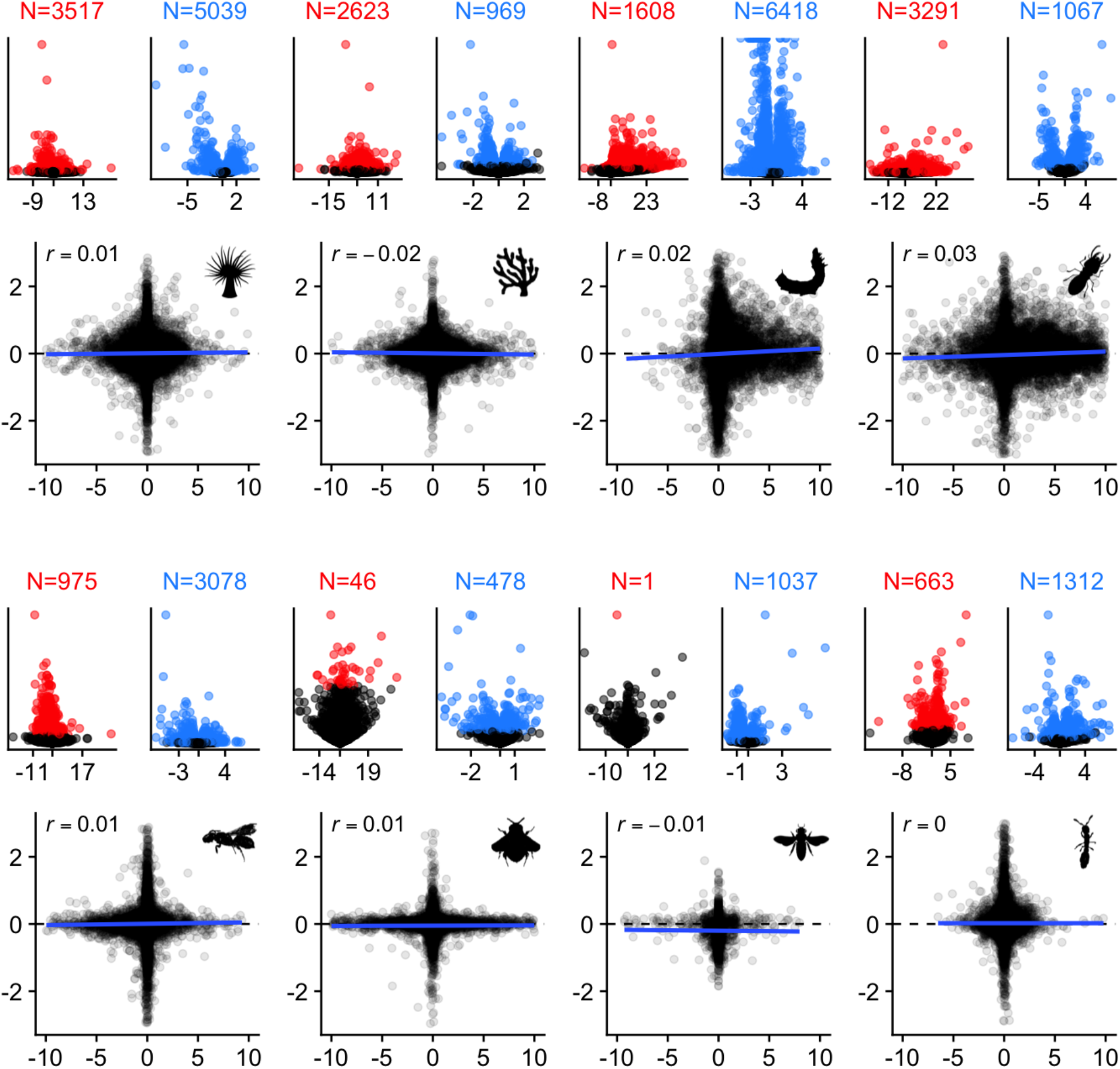
gbM and transcriptional differences between experimental conditions show little or no relationship. Each pair of volcano plots is associated with the scatterplot below. The first volcano plot shows differences in gbM, with significant genes (q-value from MethylKit < 0.1) shown in red. The second shows differences in transcription, with significant genes in blue (FDR < 0.1). The count of significant genes is given above each volcano plot. The scatterplots are the same as those shown in Figure 5, with expression differences on the Y axis and gbM differences on the X. The contrasts for each study are given in Figure 1.

## Notes

### Competing Interest Statement

The authors have declared no competing interest.

